# Targeted optimization of regulatory DNA sequences with neural editing architectures

**DOI:** 10.1101/714402

**Authors:** Anvita Gupta, Anshul Kundaje

**Affiliations:** Department of Computer Science, Stanford University; Department of Genetics, Stanford University

## Abstract

Targeted optimizing of existing DNA sequences for useful properties, has the potential to enable several synthetic biology applications from modifying DNA to treat genetic disorders to designing regulatory elements to fine tune context-specific gene expression. Current approaches for targeted genome editing are largely based on prior biological knowledge or ad-hoc rules. Few if any machine learning approaches exist for targeted optimization of regulatory DNA sequences.

Here, we propose a novel generative neural network architecture for targeted DNA sequence editing – the EDA architecture – consisting of an encoder, decoder, and analyzer. We showcase the use of EDA to optimize regulatory DNA sequences to bind to the transcription factor SPI1. Compared to other state-of-the-art approaches such as a textual variational autoencoder and rule-based editing, EDA significantly improves predicted binding of SPI1 of genomic sequences with the minimal set of edits. We also use EDA to design regulatory elements with optimized grammars of CREB1 binding sites that can tune reporter expression levels as measured by massively parallel reporter assays (MPRA). We analyze the properties of the binding sites in the edited sequences and find patterns that are consistent with previously reported grammatical rules which tie gene expression to CRE binding site density, spacing and affinity.

## 1 Introduction

Recent generative models for genomic DNA sequences, such as generative adversarial networks, variational autoencoders, and recurrent neural networks, have largely focused on ab initio generation of biological sequences from distributions learned over a large collection of exemplar sequences[10, 6, 7]. However, generative models have been shown to suffer from low diversity – falling into the failure mode of producing generic sequences with high likelihood [8]. Generative models that are capable of editing an existing sequence, rather than generating an entirely new sequence from scratch, may be able to draw from the natural diversity present in biological sequences, while still allowing useful changes to the data. Also, many genome engineering applications typically require editing an existing DNA sequence in order to knock out or repair disease genes or modify regulatory DNA to modulate gene expression in specific cell types and states.

Machine learning approaches for editing existing sequences for desired properties have been significantly less well studied than ab-initio generative models. Guu *et al* proposed a neural editor for natural language to transform an input sentence into an output based on a sampled edit vector; however, the edit vectors are latent and must be interpreted after training [8].

Here, we propose a novel Encoder-Decoder-Analyzer (EDA) neural network architecture that radically departs from status quo methods [8]. EDA combines recurrent sequence-to-sequence models, latent vectors based on an explicit predictor of desired sequence properties and adversarial example generation techniques. EDA automatically generates candidate modifications to prespecified regulatory DNA sequences to optimize specific properties such as binding of transcription factors or reporter gene expression. This model represents a unique approach to edit sequences for desired properties by leveraging existing supervised learning models that can accurately map regulatory sequences to specific properties. We showcase the EDA model on two pilot case studies. In the first case study, we use EDA to edit regulatory DNA sequences to increase the binding probability of a transcription factor SPI1 by leveraging *in vivo* genome-wide binding profiles (ChIP-seq) for SPI1. In the second case study, we use EDA to generate candidate regulatory sequences containing configurations of binding sites of the CREB1 transcription factor that can produce a desired gene expression readout as measured by a massively parallel reporter assay (MRPA) from Davis *et al,* 2019 [3]. The EDA approach significantly outperforms existing state-of-the-art approaches.

## 2 Methods

### Sequence Variational Autoencoder (SVAE) as a baseline method

The sequence variational autoencoder (SVAE) for editing is based off the recurrent architecture described in [1]; the encoder produces the parameters (*μ*, Σ) of a Gaussian distribution in latent space, from which *z*, a latent vector encoding the sequence *x*, is sampled. The decoder attempts to reconstruct the input sequence from *z*. The loss function of the VAE is given in Equation 1. For editing, the latent space of the SVAE was perturbed through the addition of Gaussian Noise 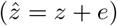 where 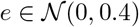 as proposed in Guu *et al* [8].

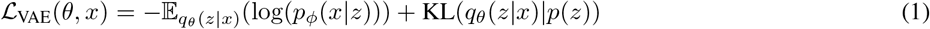

### Encoder Decoder Analyzer (EDA) Custom Architecture

Our novel architecture called EDA consists of three deep neural network components: an Encoder, Decoder, and Analyzer. The Encoder and Decoder are recurrent neural networks (RNNs) with attention. The encoder architecture used here consisted of an embedding layer followed by a recurrent layer. The embedding layer contained learnable weights and output size *h* = 256, and the embedded outputs are then fed into a one layer bidirectional GRU, with dropout of 0.1. Similarly, the decoder consists of an embedding layer (output size *h* = 256 and dropout *p* = 0.1) which creates an embedding for the input base pair at each time step, followed by attention over the encoder outputs. The outputs from the attention layer are concatenated with the input at each time step, and fed into a dense connected layer with a ReLU activation function. The outputs from this dense layer are fed into a one layer GRU with dropout *p* = 0.1, followed by a fully connected layer with a softmax activation function. The output from the decoder at each time step is the predicted next token in the sequence. Conceptually, the Encoder learns to transform any one-hot encoded input DNA sequence to a compact latent representation. The Decoder learns to generate an output DNA sequence given a specific instantiation of the latent representation learned by the encoder. The Analyzer is a neural network that can map the latent representation of a DNA sequence to a specific property that we are typically interested in optimizing. Here, we use a convolutional neural network (CNN) as the Analyzer architecture, although it can be any differentiable architecture. Details of the Analyzer architecture are as follows: the model consists of three convolutional layers (15 filters, kernel size of 3, padding of 1), an average pooling layer, and two densely connected layers. The activation function following each layer was a ReLU activation, save for the last layer, which had a sigmoid activation function in the classification setting, and no nonlinearity in the regression setting.

The procedure for editing in the EDA architecture consists of three phases.

**Stage 1: Training Encoder-Decoder**. The Encoder-Decoder seq2seq architecture is trained as an autoencoder, with loss equal to the categorical cross entropy between the softmax outputs and one-hot-encoded next base pair, summed over every position in the sequence. The loss was minimized with the Adam optimizer with learning rate 0.001. The Encoder takes in sequence *x* and produces a latent space embedding *z*, while the Decoder takes *z* and attempts to reproduce the original sequence *x*.
**Stage 2: Training Analyzer** The same Encoder from stage 1 is also followed by an Analyzer module, which takes in the latent state *z* of a sequence from the Encoder, and produces an output prediction *y* of a property of the sequence.
**Stage 3: Editing**. Given input sentence *x*, the encoder produces the latent state embedding *z* for the sentence. We update the latent state to minimize the loss function 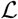(the binary cross entropy loss) between the analyzer’s prediction 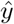 and the desired score *y* via the Fast Gradient Sign Method (FGSM) [5] as in Equation 2.

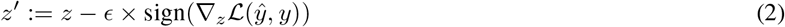

#### Algorithm 1 EDA Architecture Editing.

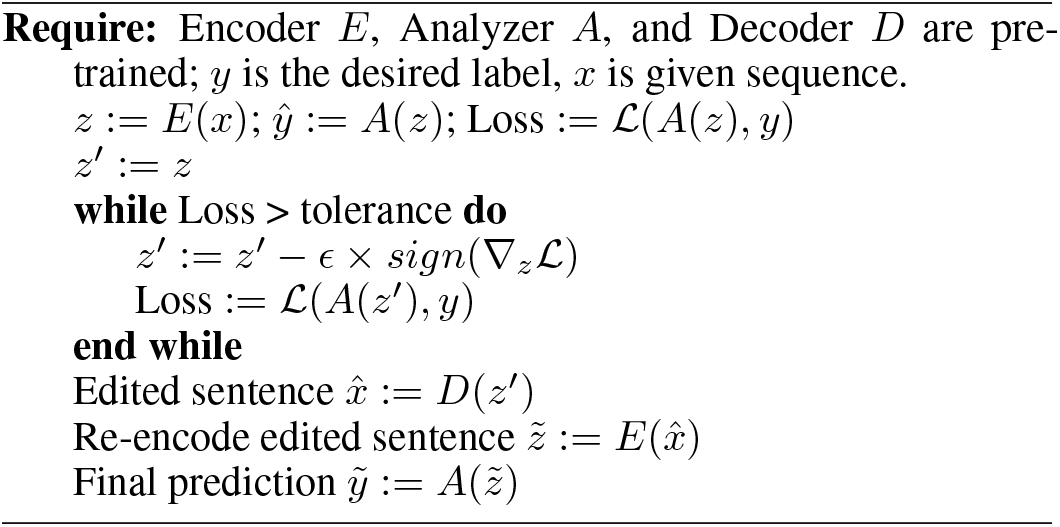

The latent state is updated until the loss is approximately 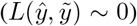. The decoder produces the edited sequence 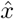 from the modified latent representation *z*′. Epsilon (*ϵ*) is a hyperparameter varying between zero and one corresponding to the size of steps taken in the latent space.

The pseudocode for the EDA training is shown in Algorithm 1. The training algorithm includes an additional step where the desired property (e.g. SPI binding or reporter expression) is predicted from the edited sentence 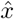. This step is necessary as the latent representation 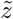 of the decoded sequence 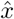 may not be the same as the perturbed latent representation *z*′ due to noise in the decoding process.

## 3 Optimizing regulatory DNA sequences for binding of the SPI1 transcription factor

### Dataset

43,787 reproducibly-identified peaks from an ENCODE ChIP-seq experiment targeting the SPI1 transcription factor in lymphoblastoid cell line GM12878 (GEO GSM803531) were used as the positive labeled set of putative SPI1 bound sequences. The negative labeled set was constructed from an equal number of non-overlapping unbound 200 bp sequences from the human genome. Datapoints from chromosomes 1 and 2 were used as the test and validation set, respectively.

### Baselines

An SVAE model was trained as a neural baseline, as described above. A simple rule-based editing model was also constructed that randomly adds the SPI1 consensus binding site (“AGGAA”) if not already present in the sequence.

### Evaluation Method

Edited sequences are evaluated on three quantitative metrics: similarity to the original DNA sequence, predicted binding score (probability) of SPI1, and percent of sequences with matches to known SPI1 binding motifs [9]. Binding score is measured by an independent CNN model trained to discriminate 200bp sequence labeled as bound by SPI1 ChIP-seq data from a balanced number of background sequences from the geneome. This independent CNN model achieves AUROC of 0.979 and AUPRC of 0.978 on a held-out test set, where, for training the independent model, datapoints from chromosomes 1 and 2 were used as the test and validation set, specifically, while all other sequences were used in the training set. Similarity of edited sequences to the original sequences was calculated by the gapped kmer-mismatch (GKM) kernel, which evaluates DNA sequence similarity based on gapped kernel overlap [4]. We also used the BLEU-4 score, a metric more commonly used in NLP translation as another measure of sequence similarity.

### 3.1 SPI1 Editing Results

#### Training Results

The loss curve for the SVAE, as well as the Encoder-Decoder portion of the EDA Architecture is shown in Figure 1. Whereas the SVAE loss is extremely noisy and difficult to optimize during training, The Encoder-Decoder of EDA achieves average edit distance of 8.8 out of a maximum possible edit distance of 150 between the input and output sequences after 20,000 iterations of training, which shows that the autoencoder is accurately learning to replicate the sequence. The accuracy of the analyzer in the EDA architecture is 92.674%, with AUROC of 0.979 and AUPRC of 0.978.

**Figure 1:**
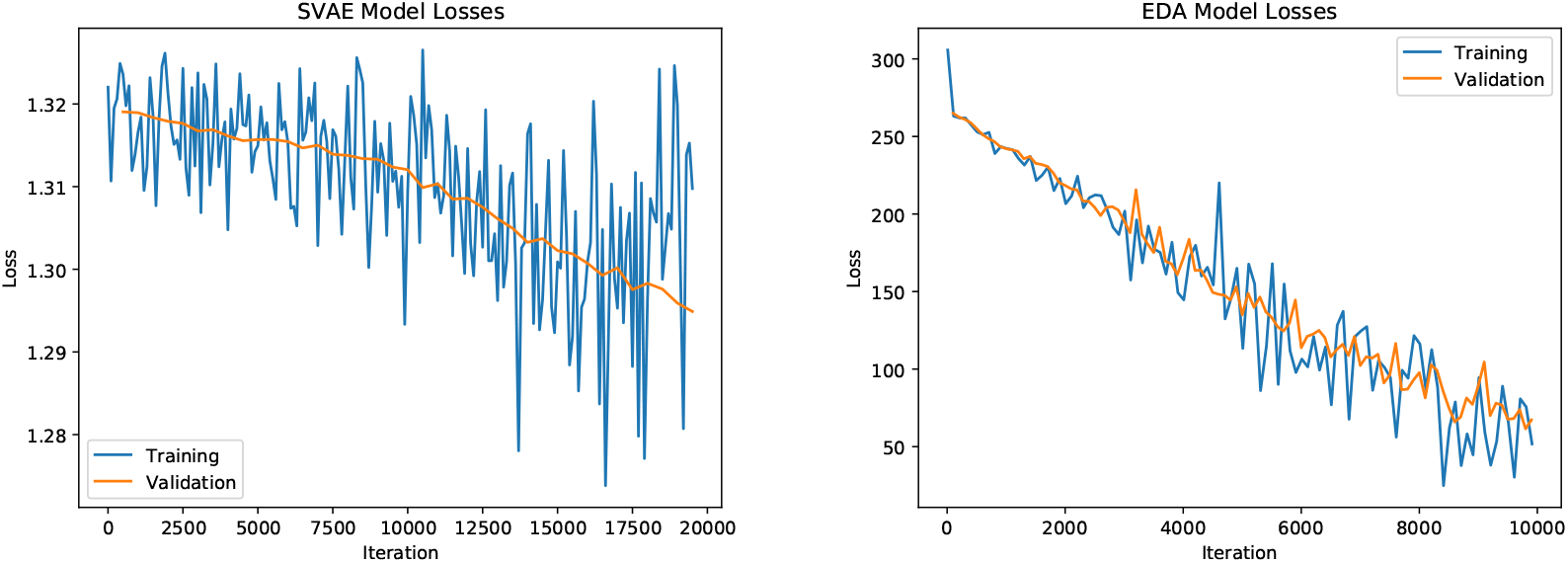
Training and validation loss curves of EDA Architecture (left) and VAE architecture (right).

#### EDA Edited Sequences

500 randomly selected sequences from the balanced test set were edited using EDA, where previously bound sequences in the test set were classified based on overlap with SPI1 ChIP-seq peaks). Results are shown in Table 1. Edits from the EDA architecture demonstrate high similarity (52.39% on average) to the original sequences, whereas the SVAE edits achieve similarity of only 5.2%. Any two random sequences from the set have GKM similarity of 2.048%. Thus, rather than editing, the SVAE appears to be sampling separate sequences.

**Table 1:** Comparison of Editing Methods in terms of sequence similarity (out of 1), BLEU-4 score (out of 100), and percentage of sentences predicted positive.

|  | GKM Similarity | BLEU-4 | % Predicted Positive | % Binding Site |
| --- | --- | --- | --- | --- |
| EDA $\epsilon = 0.55$ | **0.524** | **86.8** | 73.4 | 31.8 |
| EDA $\epsilon = 0.99$ | 0.491 | 85.1 | **84.4** | **34.2** |
| VAE | 0.052 | 33.12 | 16.0 | 20.6 |
| Rule Baseline | 0.965 | 98.3 | 45.0 | 100 |

Overall, 84.4% of EDA-edited sequences are predicted to bind to SPI1 by the independent CNN model trained on SPI1 ChIP-seq data. 34.2% of these sequences contain a deterministic match to the SPI1 motif “AGGAA”, and 63.4% of sequences contain the “GGAA” portion, which is has the highest information content in the SPI1 PWM [9]. The rule-based baseline achieves only 45% sequences predicted positive, similar to the probability that any randomly chosen test set sequence would be predicted positive. Thus, editing these sequences to optimize for binding score is more complex than simply inserting high affinity SPI1 binding sites.

**Table 2** shows a DNA sequence which initially had a low binding score on the independent model, whose edit received a high score. The EDA model modifies the area in the initial sequence in gray into the full SPI1 binding motif shown in orange.

**Table 2:**
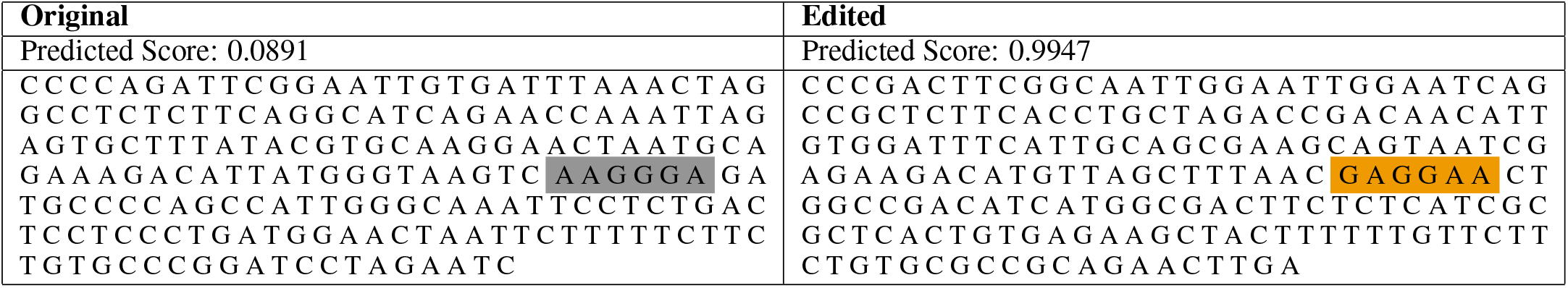
Original sequence and edited sequence from the EDA architecture. The SPI1 motif is highlighted in orange.

## 4 Optimizing Reporter Expression of regulatory DNA sequences containing CREB1 binding site grammars

### Dataset

The CRE MPRA dataset from Davis et al. measures reporter gene expression of a library of DNA sequences with various configurations of CREB1 binding sites by varying motif strength, density, spacing, and distance from the core promoter[3]. The genomic MPRA dataset consists of 3480 sequences of 150bp within 3 backgrounds with different combinations and locations of CREB1 binding sites. Davis *et al.* define a strong CREB1 consensus binding site as “TGACGTCA”, and a weak binding site is “TGAAGTCA”. Reporter expression of the library is measured by the log ratio of counts of RNA barcode reads of a sequence to the count of DNA reads. A histogram of log fold change expression levels is shown in Figure 2. Seventy percent of this dataset was randomly selected for training, with twenty percent for validation and ten percent for testing.

**Figure 2:**
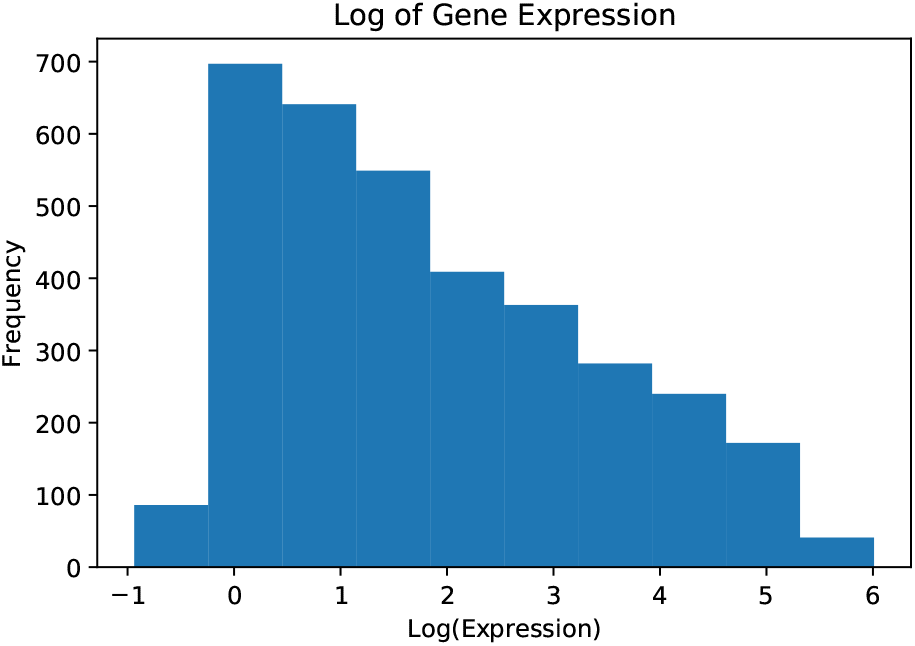
Histogram of log(expression) levels for sequences in CREB1 MPRA dataset; expression ranges widely, from less than zero, meaning that number of RNA barcoded reads are less than the original DNA reads, to six.

From analysis of the MPRA dataset, Davis *et al* find four main correlations between CREB1 binding site configurations in the library and corresponding reporter expression levels: 1) number of strong CRE binding sites positively correlates with expression, 2) weak binding sites increase expression given the presence of at least one strong binding site, 3) higher expression occurs with shorter distance of CRE binding sites to the core promoter, and 4) spacing between CRE binding sites modulates periodicity of expression, as two strong binding sites are moved along the sequence.

Here, the editing task is to optimize sequences for particular expression profiles; in particular, to edit MPRA sequences that have high measured reporter expression (log(expression)) ≥ 5) to new sequences that have low predicted expression, and vice versa. As several correlations between sequence properties and expression are already discussed by Davis *et al.,* we investigate whether the edited sequences show evidence of these previously discovered rules.

### Independent Analyzer

As an independent predictor from the analyzer in the EDA architecture, we train a log-linear model of expression levels similar to Davis *et al.,* with expression predicted from the number of strong and weak binding sites, sequence background, average spacing between CREB1 sites, and distance from the minimal promoter element; in addition, polynomial features of degree 2 were used to model interaction terms. This simple model achieves *R*^2^ = 0.801 on a held out test set consisting of 10 percent of the training data, which was the same test set as used for the training of the EDA architecture (**Figure 3a.**). This independent log linear model was used to evaluate the edited sequences from the EDA model.

**Figure 3:**
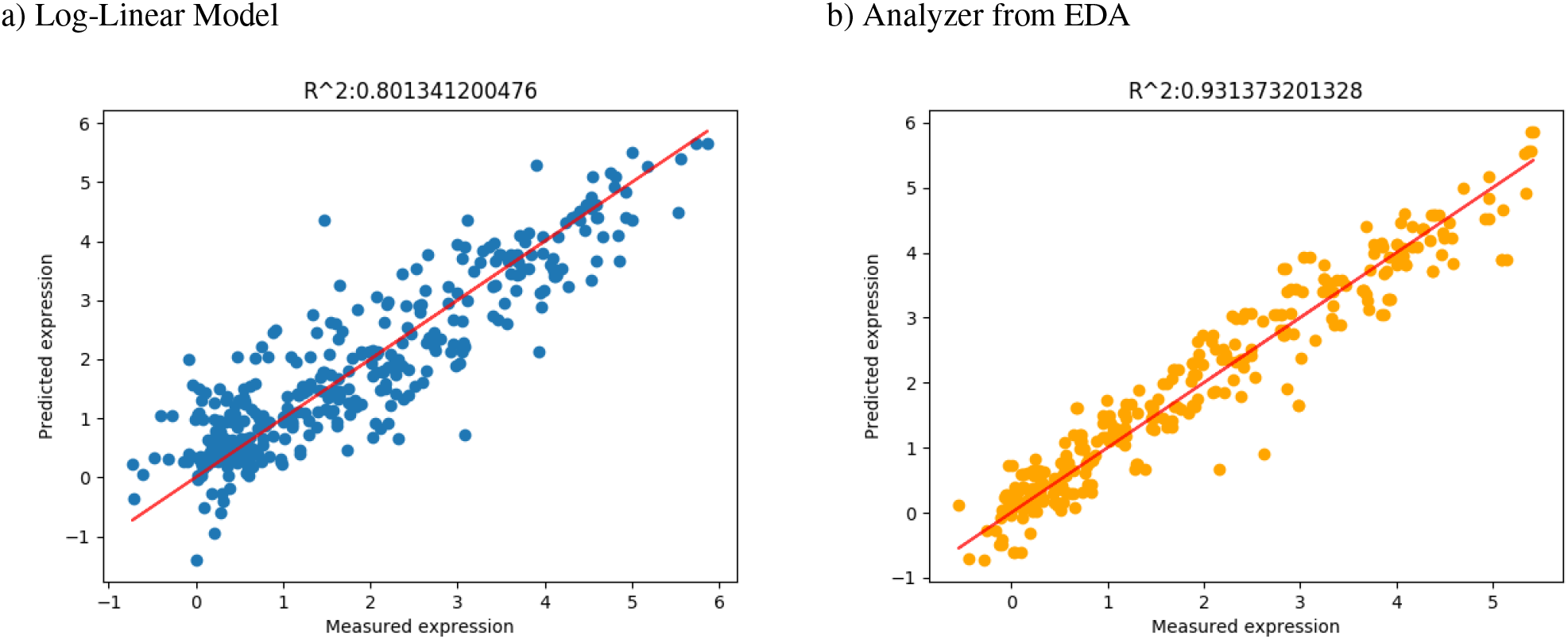
Predicted expression versus measured expression for a) independent log-linear model, and b) analyzer from EDA architecture

### Training Results

The components of the EDA model were trained on the CRE MPRA dataset to learn a latent representation of the sequences through the encoder-decoder portion, and to predict reporter expression levels from this latent representation through the analyzer. As described in Section 2, the encoder and decoder were both recurrent neural networks, where the decoder has softmax attention over the encoder outputs. The seq2seq autoencoder achieved an average edit distance of 6.06 between input and output after training for 10,000 iterations. The analyzer, as above, was a CNN architecture with three convolutional layers, each followed by a ReLU activation, average pooling, and two dense layers; this architecture achieved *R*^2^ = 0.93 on a held out test set; the analyzer’s predicted expressions correlate well with measured expressions, as shown in **Figure 3b**.

### EDA Editing Results

Here, our editing task is to edit CREB1 MPRA sequences from a held out test set that exhibit high measured expression to new sequences with low expression and vice versa. We further evaluate whether the resulting edited sequences displayed known patterns of CRE binding site placement elucidated by Davis *et al*. 204 sequences with measured MPRA log(expression) <= 0 were edited using EDA to a higher desired target level of log(expression) = 5.0. 96 sequences with measured MPRA log(expression) >= 5 were edited using EDA to obtain a lower target level of log(expression) = 0.

As evaluated by the independent analyzer, 79.4% of sequences to be edited from low expression to high expression were predicted to have higher expression post editing. 100% of sequences edited from high to low expression were predicted to have lower expression post editing. The histograms of log expression levels both before and after editing are shown in **Figure 4**.

**Figure 4:**
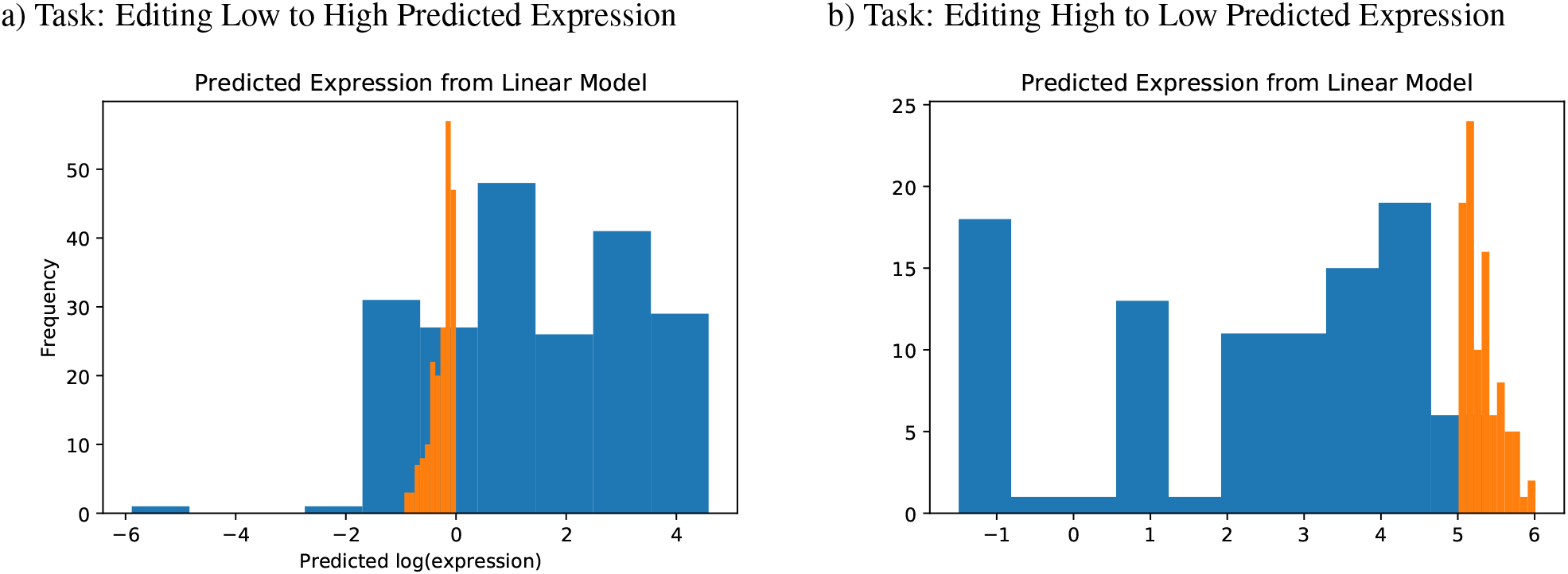
Histograms of log expression levels before and after editing by the EDA architecture and evaluation on an independent linear model. Log expression of original sequences is shown in orange, while log expression of edited sequences is shown in blue, where expression of edited sequences is predicted by the independent model. Edits targeted from low to high expression are shown in a, and edits from high to low expresssion are shown in b. Here, low expression was defined as log(expression) ≤ 0.0, and high expression was defined as log(expression) ≥ 5.0.

Next, we inspected several representative edited sequences in order to evaluate whether they contained CREB1 binding site configurations that were previously associated with high and low expression read outs, where examples are shown in **Figure 5**.

**Figure 5:**
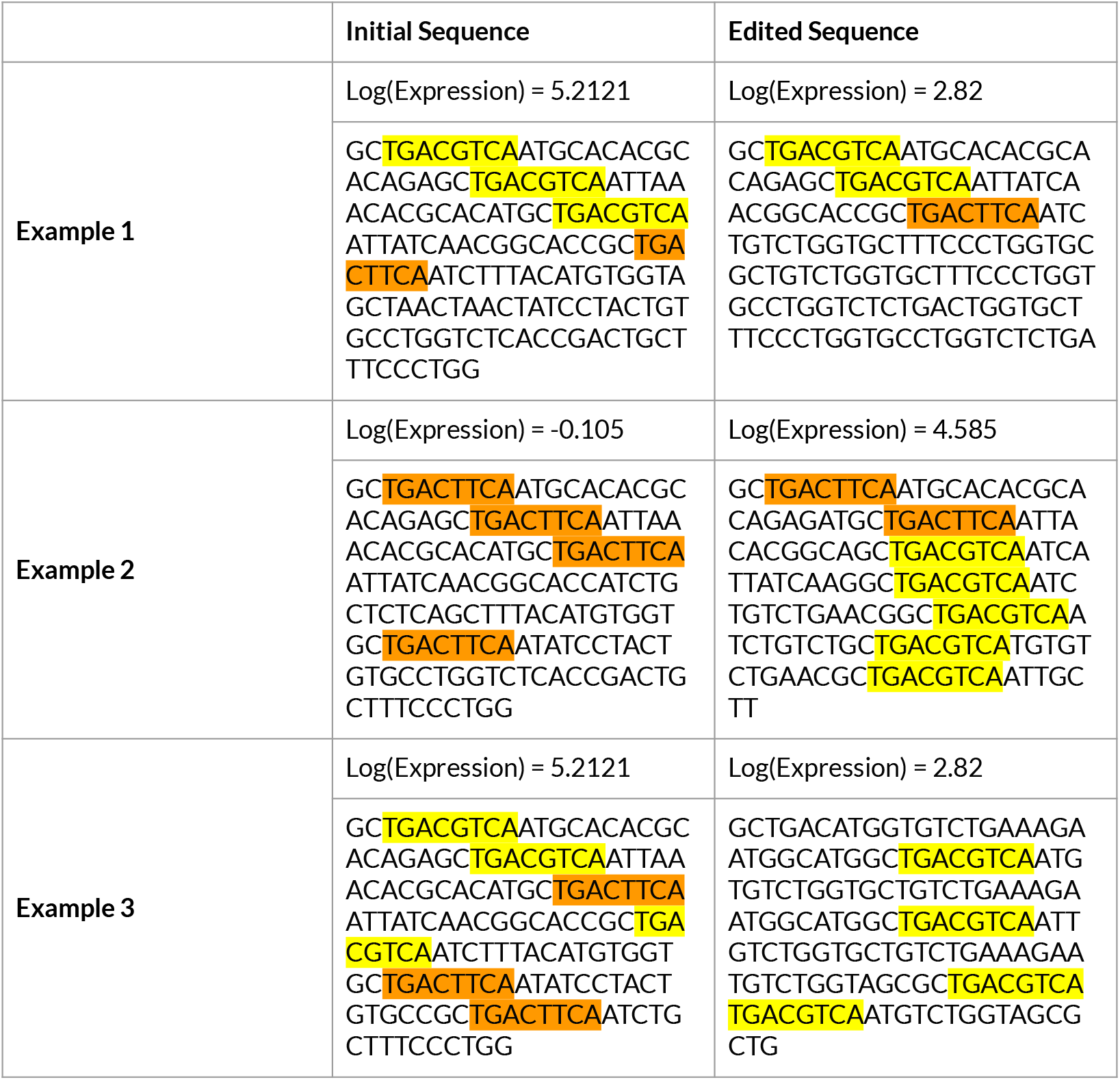
Strong CREB1 binding motif is highlighted in yellow, while weak motif is highlighted in orange. Predicted expression, measured by the log of the ratio of RNA to DNA counts, is shown above each sequence.

Example 1 illustrates the result that number of strong CREB1 binding sites is positively correlated with expression levels, as the edited sequence has the third strong CREB1 binding site changed to a weak site, resulting in expression being predicted to be more than 2x lower after editing.

In example 2, the original sequence has four weak binding sites and exhibits low measured expression (−0.101). After editing, the third weak binding site is changed to a strong binding site, the fourth weak binding site is deleted, and four additional strong binding sites are added to the sequence. This edit, with predicted expression 4.58, also displays the result found in Davis *et al*, that number of weak binding sites increases reporter gene expression given the presence of at least one strong binding site.

In the third example, EDA moves a CREB1 motif moved further away from the minimal promoter element in order to reduce expression. This transformation of the sequence aligns with the previous reported observation from the MPRA study that distance of motifs from the core promoter negatively correlated with expression. These examples are presented with the caveat that they cannot be used to show that the model has “learned” particular rules – only that the results from the EDA architecture align with known experimental correlations between CREB1 binding sites and reporter expression from the MPRA study.

## 5 Conclusion

The EDA architecture proposed for targeted genomic editing brings together a broad array of techniques from attention-based seq2seq models, adversarial example generation, and computer vision. The architecture leverages existing genomic predictors to generate candidate modifications of sequences with diverse properties such as binding probability of a transcription factor or reporter gene expression levels. In the first case study of optimizing binding of the SPI1 transcription factor, we compared the EDA model to existing neural baselines – such as the Sequence VAE model and a rule-based baseline – and showed that EDA vastly improves upon existing models in both predicted binding affinity and similarity of original to edited sequences.

In the second case study, where we optimized binding site configurations of the CREB1 transcription factor in regulatory DNA sequences to tune reporter expression levels, we showed that a high proportion of the edited sequences show the desired shift in expression as predicted by an independent model. Furthermore, several edited sequences displayed CREB1 motif configurations in terms of binding site strength, density and position that agreed with previously derived rules.

This study primarily serves as a proof of concept and introduction of a novel neural architecture for targeted DNA sequence editing. In this work, we used independent predictors of the desired properties of DNA sequences to computationally validate the edited sequences. In the near future, we plan to provide experimental validation of the properties of edited sequences as more definitive support for our approach. EDA is very flexible and can also be easily adapted to other applications involving targeted DNA and RNA editing. We expect further advances in generative models that can perform targeted editing of biological sequences such as DNA, RNA and proteins have the potential to complement and improve the precision of experimental approaches for genome engineering and synthetic biology.

## 6 Acknowledgements

We would like to thank Georgi Marinov for his help with processing and understanding the MPRA dataset. We would like to thank the authors of Davis *et al.* [3] for sharing their MPRA data pre-publication. We would like to thank Avanti Shrikumar and other members of the Kundaje lab for helpful discussions.

This work was supported by NIH grants 1DP2GM123485, 1U01HG009431 and 1R01HG00967401 awarded to AK.

